# GWAS meta-analysis (N=279,930) identifies new genes and functional links to intelligence

**DOI:** 10.1101/184853

**Authors:** Jeanne E Savage, Philip R Jansen, Sven Stringer, Kyoko Watanabe, Julien Bryois, Christiaan A de Leeuw, Mats Nagel, Swapnil Awasthi, Peter B Barr, Jonathan R I Coleman, Katrina L Grasby, Anke R Hammerschlag, Jakob Kaminski, Robert Karlsson, Eva Krapohl, Max Lam, Marianne Nygaard, Chandra A Reynolds, Joey W Trampush, Hannah Young, Delilah Zabaneh, Sara Hägg, Narelle K Hansell, Ida K Karlsson, Sten Linnarsson, Grant W Montgomery, Ana B Muñoz-Manchado, Erin B Quinlan, Gunter Schumann, Nathan Skene, Bradley T Webb, Tonya White, Dan E Arking, Deborah K Attix, Dimitrios Avramopoulos, Robert M Bilder, Panos Bitsios, Katherine E Burdick, Tyrone D Cannon, Ornit Chiba-Falek, Andrea Christoforou, Elizabeth T Cirulli, Eliza Congdon, Aiden Corvin, Gail Davies, Ian J Deary, Pamela DeRosse, Dwight Dickinson, Srdjan Djurovic, Gary Donohoe, Emily Drabant Conley, Johan G Eriksson, Thomas Espeseth, Nelson A Freimer, Stella Giakoumaki, Ina Giegling, Michael Gill, David C Glahn, Ahmad R Hariri, Alex Hatzimanolis, Matthew C Keller, Emma Knowles, Bettina Konte, Jari Lahti, Stephanie Le Hellard, Todd Lencz, David C Liewald, Edythe London, Astri J Lundervold, Anil K Malhotra, Ingrid Melle, Derek Morris, Anna C Need, William Ollier, Aarno Palotie, Antony Payton, Neil Pendleton, Russell A Poldrack, Katri Räikkönen, Ivar Reinvang, Panos Roussos, Dan Rujescu, Fred W Sabb, Matthew A Scult, Olav B Smeland, Nikolaos Smyrnis, John M Starr, Vidar M Steen, Nikos C Stefanis, Richard E Straub, Kjetil Sundet, Aristotle N Voineskos, Daniel R Weinberger, Elisabeth Widen, Jin Yu, Goncalo Abecasis, Ole A Andreassen, Gerome Breen, Lene Christiansen, Birgit Debrabant, Danielle M Dick, Andreas Heinz, Jens Hjerling-Leffler, M Arfan Ikram, Kenneth S Kendler, Nicholas G Martin, Sarah E Medland, Nancy L Pedersen, Robert Plomin, Tinca JC Polderman, Stephan Ripke, Sophie van der Sluis, Patrick F Sullivan, Henning Tiemeier, Scott I Vrieze, Margaret J Wright, Danielle Posthuma

**Author notes:** These authors contributed equally (Jeanne E Savage, Philip R Jansen). Correspondence to: Danielle Posthuma: Department of Complex Trait Genetics, VU University, De Boelelaan 1085, 1081 HV, Amsterdam, The Netherlands.

## Abstract

Intelligence is highly heritable^1^ and a major determinant of human health and well-being^2^. Recent genome-wide meta-analyses have identified 24 genomic loci linked to intelligence^3–7^, but much about its genetic underpinnings remains to be discovered. Here, we present the largest genetic association study of intelligence to date (N=279,930), identifying 206 genomic loci (191 novel) and implicating 1,041 genes (963 novel) via positional mapping, expression quantitative trait locus (eQTL) mapping, chromatin interaction mapping, and gene-based association analysis. We find enrichment of genetic effects in conserved and coding regions and identify 89 nonsynonymous exonic variants. Associated genes are strongly expressed in the brain and specifically in striatal medium spiny neurons and cortical and hippocampal pyramidal neurons. Gene-set analyses implicate pathways related to neurogenesis, neuron differentiation and synaptic structure. We confirm previous strong genetic correlations with several neuropsychiatric disorders, and Mendelian Randomization results suggest protective effects of intelligence for Alzheimer’s dementia and ADHD, and bidirectional causation with strong pleiotropy for schizophrenia. These results are a major step forward in understanding the neurobiology of intelligence as well as genetically associated neuropsychiatric traits.

---

We performed a genome wide meta-analysis of 16 independent cohorts totaling 279,930 participants of European ancestry and 9,398,186 genetic variants passing quality control (**Online Methods; Supplementary Table 1; Supplementary Figure 1**). All genome-wide analyses were corrected for cohort-specific ancestry and covariates (**Supplementary Information**). Various measures of intelligence were used in each study, yet genetic correlations between cohorts (*r_g_*, **Online Methods**), were considerable (mean=0.63), warranting meta-analysis (**Supplementary Table 2; Supplementary Results 2.1**). Separate meta-analyses for children, young adults, and adults (**Online Methods**) indicated high genetic correlations between age groups (*r_g_*>0.62), and comparable single nucleotide polymorphism (SNP)-based heritability across age (*h*^*2*^*_SNP_*=0.19-0.22) (**Supplementary Table 3; Supplementary Results 2.2**). The inflation factor of the full meta-analysis was λ_GC_=1.95 (**Supplementary Table 4; Supplementary Figure 2**), with *h*^*2*^*SNP*=0.18 (SE=0.01), in line with previous findings^4,5^, and an LD score intercept^8^ of 1.09 (SE=0.02) indicated that most of the inflation could be explained by polygenic signal and large sample size^6^.

In the meta-analysis, 12,701 variants indexed by 531 independently significant SNPs (*r*^*2*^<0.6) and 246 lead SNPs in approximate linkage equilibrium (*r*^*2*^<0.1; **Online Methods**) reached genome-wide significance (GWS; *P*<5×10^−8^) (**Figure 1a; Supplementary Table 5; Supplementary Figure 3**). These were located in 206 distinct genomic loci, 191 of which are novel associations (**Supplementary Results 2.3**). Proxy replication with the correlated phenotype of educational attainment (EA; *r_g_*=0.73) in an independent sample (**Online Methods**) indicated sign concordance for 94% of GWS SNPs (*P*<1×10^−300^) and evidence of replication for 51 loci (**Supplementary Results 2.3.2; Supplementary Table 6**). Using polygenic score prediction^9,10^ (Online Methods) we show that the current results explain up to 5.4% of the variance in four independent samples (**Supplementary Table 7, Supplementary Results 2.3.3**).

**Figure 1.**
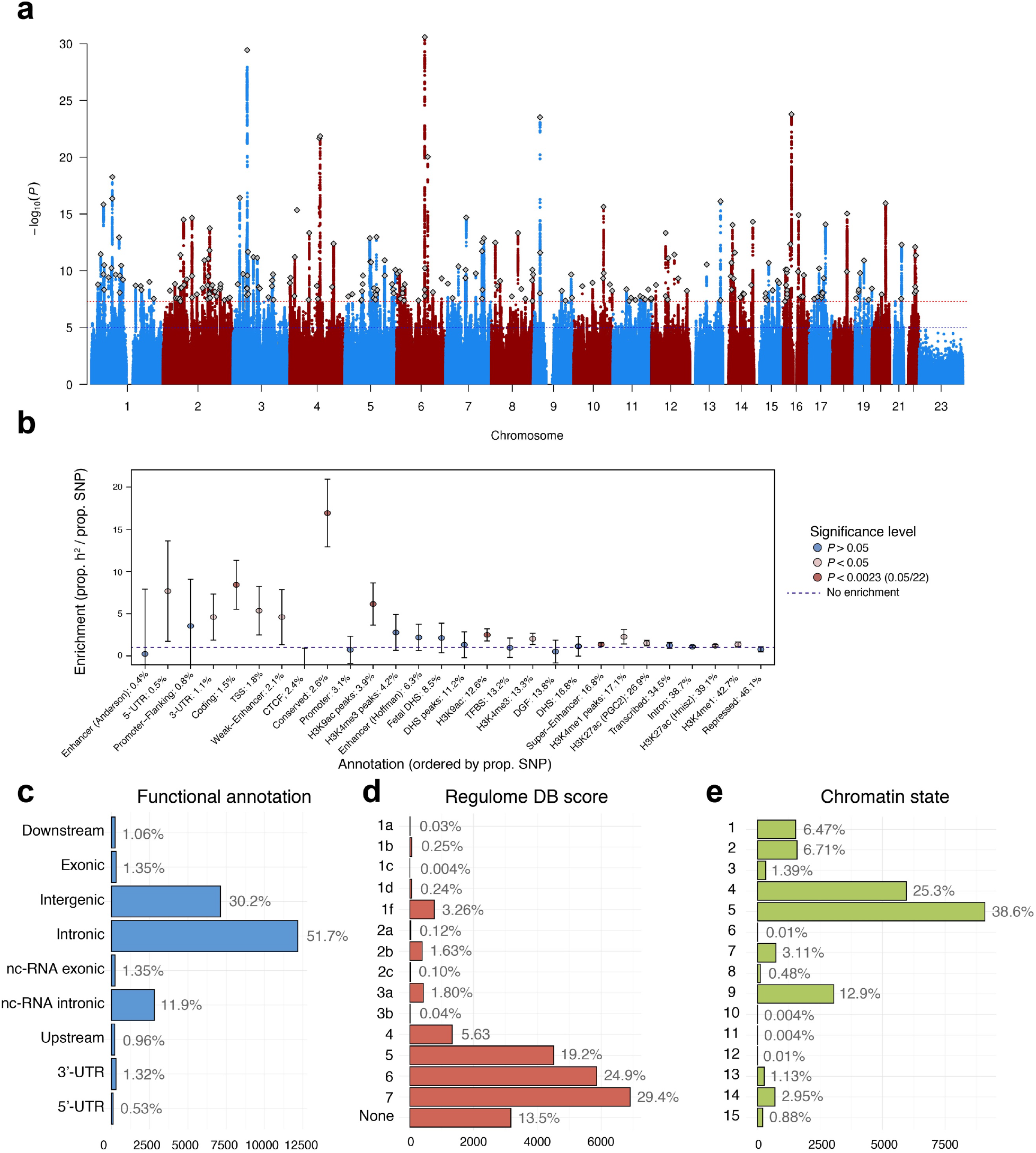
SNP-based associations with intelligence in the GWAS meta-analysis of N=279,930. (**a**) Manhattan plot showing the ─log10 transformed *P*-value of each SNP on the y-axis and base pair positions along the chromosomes on the x-axis. The dotted red line indicates genome-wide significance (*P*<5×10^−8^), the blue line the threshold for suggestive associations (*P*<1×10^−5^). Independent lead SNPs are indicated by a diamond. **(b)** Heritability enrichment of 28 functional SNP annotations calculated with stratified LD score regression TSS = Transcription Start Site; CTCF=CCCTC-binding factor; DHS=DNase Hypersensitive Site. **(c)** Distribution of functional consequences of SNPs in genomic risk loci in the meta-analysis. **(d)** Distribution of RegulomeDB score for SNPs in genomic risk loci, with a low score indicating a higher likelihood of having a regulatory function (**Online methods**). **(e)** The minimum chromatin state across 127 tissue and cell types for SNPs in genomic risk loci, with lower states indicating higher accessibility and states 1-7 referring to open chromatin states (**Online Methods**).

We observed strong enrichment for heritability (**Online Methods; Supplementary Results 2.3.4**) of SNPs located in conserved regions of the genome (*P*=1.84×10^−12^), coding regions (*P*=7.88×10^−7^), H3K9ac histone regions/peaks (*P*<6.06×10^−5^), and super-enhancers (*P*=9.61×10^−5^) (**Figure 1b; Supplementary Table 8**). Conserved regions have previously been implicated for intelligence^11^ but coding regions have not. Heritability was disproportionately found among common variants (**Supplementary Figure 5**) with greatest enrichment for SNPs with a minor allele frequency (MAF) between 0.4 and 0.5 (*P*=5.81×10^−12^), but was distributed proportionally across chromosomes (**Supplementary Figure 6**).

Functional annotation of all SNPs (*n=*23,552) in the associated loci was performed using FUMA^12^ (**Online Methods**). SNPs were mostly located in intronic (*n*=12,171; 51.7%) and intergenic areas (n=9,923; 42.1%) (**Supplementary Table 9; Figure 1c**), yet 6.3% (1,473 SNPs) were annotated to functional genic regions, with 1.4% (318 SNPs) being exonic. Of these, 89 (41 GWS) SNPs were exonic non-synonymous (ExNS) (**Table 1, Supplementary Results 2.3.5**). Convergent evidence of strong association (*Z*=9.74) and the highest observed probability of a deleterious protein effect (CADD^13^ score=34) was found for rs13107325. This missense mutation (MAF=0.065) in *SLC39A8* was the lead SNP in locus 71 and the ancestral allele C was associated with higher intelligence scores. The effect sizes for ExNS were individually small, with each effect allele 0.01 to 0.05 standard deviations. **Table 1, Supplementary Table 9** and **Supplementary Results 2.3.5** present a detailed catalog of the functional impact of variants in the genomic risk loci. Apart from protein consequences, the implicated SNPs also showed some evidence of indirect functional effects: 4.4% had a RegulomeDB^14^ score of 1a-1f (**Figure 1d**), suggesting a regulatory function, and the majority of SNPs (81.6%) were in open chromatin regions^15,16^, as indicated by a minimum chromatin state of 1-7 (**Figure 1e**).

**Table 1.** Exonic non-synonymous (ExNS) variants in the genomic loci associated with intelligence and in LD (r^2^>0.6) with one of the independent GWS SNPs.

| SNP | Gene | Exon | CADD | RDB | A1 | A2 | MAF | P | Z | Effect Size |
| --- | --- | --- | --- | --- | --- | --- | --- | --- | --- | --- |
| rs4359027 | FOXO6 | 2 | 13.18 | 2b | A | G | 0.47 | 1.2E-07 | 5.29 | 0.014 |
| rs11264743 | CREB3L4 | 3 | 24.4 | 4 | T | C | 0.29 | 5.2E-07 | -5.02 | -0.015 |
| rs11264875 | NUP210L | 32 | 28.6 | 6 | T | C | 0.28 | 2.5E-06 | -4.71 | -0.014 |
| rs2232819 | METTL13 | 3 | 22.8 | 5 | A | G | 0.19 | 4.1E-07 | 5.06 | 0.017 |
| rs2820312 | LMOD1 | 2 | 2.65 | 5 | A | G | 0.34 | 1.7E-08 | -5.64 | -0.016 |
| rs1804020 | ZNF638 | 5 | 7.18 | 7 | A | G | 0.24 | 4.7E-11 | -6.58 | -0.021 |
| rs11542286 | ZNF638 | 7 | 22.4 | - | T | C | 0.09 | 2.0E-07 | -5.20 | -0.024 |
| rs3813227 | ALMS1 | 5 | 0.09 | 6 | T | C | 0.24 | 3.7E-07 | -5.08 | -0.016 |
| rs6546837 | ALMS1 | 8 | 0 | 6 | C | G | 0.24 | 4.0E-07 | -5.07 | -0.016 |
| rs6724782 | ALMS1 | 8 | 0.01 | 7 | A | T | 0.24 | 5.2E-07 | -5.02 | -0.016 |
| rs6546839 | ALMS1 | 8 | 17.18 | 7 | C | G | 0.24 | 3.7E-07 | -5.08 | -0.016 |
| rs2056486 | ALMS1 | 10 | 0.29 | 5 | T | G | 0.24 | 2.0E-07 | -5.20 | -0.016 |
| rs3828193 | CHST10 | 2 | 13.45 | - | T | G | 0.49 | 8.8E-13 | 7.15 | 0.019 |
| rs2116665 | GPD2 | 3 | 11.61 | - | A | G | 0.3 | 4.2E-09 | 5.88 | 0.017 |
| rs1064213 | PLCL1 | 2 | 26.1 | 6 | A | G | 0.48 | 8.4E-07 | -4.93 | -0.013 |
| rs9834639 | TMEM89 | 1 | 13.65 | 1f | T | G | 0.07 | 2.0E-06 | 4.76 | 0.025 |
| rs13324142 | SLC26A6 | 2 | 25.6 | 1f | T | C | 0.1 | 6.7E-08 | 5.40 | 0.024 |
| rs34759087 | LAMB2 | 22 | 23.1 | 5 | T | C | 0.12 | 3.1E-08 | 5.53 | 0.023 |
| rs13068038 | CCDC36 | 10 | 4.12 | 7 | A | C | 0.11 | 6.9E-13 | 7.18 | 0.031 |
| rs13077498 | C3orf62 | 1 | 1.48 | 4 | T | C | 0.11 | 3.2E-13 | 7.29 | 0.031 |
| rs34762726 | BSN | 5 | 0.66 | 5 | A | G | 0.29 | 3.0E-25 | 10.38 | 0.031 |
| rs2005557 | BSN | 8 | 11.52 | 2b | A | G | 0.47 | 2.6E-08 | -5.57 | -0.015 |
| rs3197999 | MST1 | 18 | 25.5 | - | A | G | 0.29 | 1.6E-23 | 10.00 | 0.029 |
| rs34823813 | RNF123 | 24 | 18.51 | 7 | A | G | 0.1 | 3.6E-12 | 6.95 | 0.031 |
| rs73079003 | CDHR4 | 8 | 5.09 | 3a | A | G | 0.13 | 3.4E-08 | 5.52 | 0.022 |
| rs1046956 | SEMA3F | 13 | 6.01 | 4 | A | T | 0.34 | 1.7E-09 | 6.02 | 0.017 |
| rs11177 | GNL3 | 3 | 22.9 | - | A | G | 0.38 | 5.7E-11 | 6.55 | 0.018 |
| rs2289247 | GNL3 | 11 | 12.82 | - | A | G | 0.41 | 1.4E-10 | 6.42 | 0.017 |
| rs1029871 | NEK4 | 5 | 24.1 | 1f | C | G | 0.38 | 5.3E-10 | 6.21 | 0.019 |
| rs3617 | ITIH3 | 9 | 0.16 | - | A | C | 0.45 | 4.6E-08 | 5.47 | 0.015 |
| rs61739170 | DHFRL1 | 2 | 0.01 | 6 | C | G | 0.26 | 5.8E-06 | -4.54 | -0.014 |
| rs2269495 | TNIP2 | 6 | 10.93 | - | A | G | 0.44 | 7.3E-07 | -4.95 | -0.013 |
| rs3795243 | NCAPG | 4 | 10.85 | 5 | C | G | 0.13 | 4.3E-11 | -6.59 | -0.026 |
| rs34811474 | ANAPC4 | 20 | 23.8 | 6 | A | G | 0.23 | 4.5E-16 | 8.12 | 0.027 |
| rs13107325 | SLC39A8 | 8 | 34 | 5 | T | C | 0.08 | 2.1E-22 | -9.74 | -0.048 |
| rs17610219 | TTC29 | 6 | 1.91 | 7 | A | G | 0.38 | 4.2E-08 | 5.48 | 0.015 |
| rs2240695 | PCDHA1 | 1 | 21.9 | - | T | G | 0.47 | 3.5E-08 | 5.52 | 0.015 |
| rs9686540 | PCDHA2 | 1 | 2.31 | 5 | A | G | 0.47 | 8.6E-08 | 5.35 | 0.014 |
| rs7701755 | PCDHA3 | 1 | 22.6 | 4 | T | G | 0.47 | 1.2E-07 | 5.29 | 0.014 |
| rs2240694 | PCDHA3 | 1 | 23 | - | A | G | 0.47 | 5.1E-08 | 5.45 | 0.015 |
| rs3822346 | PCDHA4 | 1 | 0.02 | - | T | C | 0.47 | 4.9E-08 | 5.45 | 0.015 |
| rs4141841 | PCDHA5 | 1 | 17 | - | T | C | 0.47 | 3.3E-08 | 5.53 | 0.015 |
| rs10067182 | PCDHA7 | 1 | 0.07 | 5 | A | G | 0.47 | 5.6E-08 | 5.43 | 0.015 |
| rs41266839 | BTN3A1 | 5 | 0 | 4 | C | G | 0.11 | 1.9E-07 | 5.21 | 0.022 |
| rs13195401 | BTN2A1 | 4 | 24.4 | 5 | T | G | 0.11 | 3.3E-07 | 5.11 | 0.022 |
| rs13195402 | BTN2A1 | 4 | 24.6 | 5 | T | G | 0.11 | 1.4E-06 | 4.82 | 0.021 |
| rs13195509 | BTN2A1 | 4 | 23.6 | 1f | A | G | 0.12 | 7.9E-08 | 5.37 | 0.022 |
| rs3734542 | BTN2A1 | 8 | 7.52 | 5 | A | G | 0.12 | 1.4E-07 | 5.27 | 0.022 |
| rs3734543 | BTN2A1 | 8 | 6.66 | 5 | C | G | 0.12 | 5.4E-07 | 5.01 | 0.021 |
| rs35555795 | BTN1A1 | 7 | 0.03 | 7 | T | C | 0.12 | 1.1E-07 | 5.31 | 0.022 |
| rs3749971 | OR12D3 | 1 | 1.61 | - | A | G | 0.12 | 4.0E-09 | 5.89 | 0.025 |
| rs3735478 | ZMIZ2 | 9 | 22.7 | 1f | T | G | 0.29 | 4.2E-11 | 6.60 | 0.020 |
| rs1801195 | WRN | 26 | 0.08 | 5 | T | G | 0.46 | 1.2E-07 | 5.30 | 0.014 |
| rs79460462 | TSNARE1 | 3 | 15.14 | 5 | T | C | 0.02 | 7.4E-07 | 4.95 | 0.049 |
| rs1063739 | GPT | 1 | 0.01 | 4 | A | C | 0.47 | 2.0E-09 | -6.00 | -0.016 |
| rs4251691 | RECQL4 | 18 | 19.04 | - | T | C | 0.46 | 3.3E-08 | -5.52 | -0.015 |
| rs184457 | IER5L | 1 | 25 | 4 | A | G | 0.31 | 8.1E-10 | 6.14 | 0.018 |
| rs11191274 | GBF1 | 38 | 0.01 | 5 | A | G | 0.13 | 1.8E-06 | -4.78 | -0.019 |
| rs34473884 | PPP2R2D | 6 | 24.6 | 5 | A | G | 0.25 | 5.5E-08 | 5.43 | 0.017 |
| rs2030166 | NDUFS3 | 5 | 3.13 | 6 | T | C | 0.35 | 1.3E-08 | -5.68 | -0.016 |
| rs1064608 | MTCH2 | 12 | 25.4 | 6 | C | G | 0.35 | 1.1E-08 | -5.72 | -0.016 |
| rs4926 | SERPING1 | 8 | 23.5 | 5 | A | G | 0.27 | 1.6E-07 | -5.25 | -0.016 |
| rs55865069 | KMT2D | 4 | 19.8 | 5 | T | C | 0.03 | 1.9E-06 | 4.77 | 0.038 |
| rs4647899 | AKAP6 | 13 | 13.13 | 5 | A | T | 0.3 | 2.4E-07 | 5.17 | 0.015 |
| rs17524906 | DMXL2 | 11 | 12.26 | 7 | A | G | 0.24 | 4.6E-07 | 5.04 | 0.016 |
| rs16973457 | FAM154B | 4 | 27.7 | 6 | T | C | 0.49 | 2.1E-07 | 5.19 | 0.014 |
| rs7140 | SPNS1 | 11 | 16.6 | - | A | C | 0.3 | 3.4E-09 | 5.91 | 0.017 |
| rs12949256 | ARHGAP27 | 5 | 11.97 | 4 | T | C | 0.19 | 2.8E-07 | -5.14 | -0.018 |
| rs16940674 | CRHR1 | 6 | 12.86 | 1f | T | C | 0.23 | 1.2E-07 | -5.29 | -0.017 |
| rs16940681 | CRHR1 | 13 | 1.76 | 4 | C | G | 0.23 | 3.5E-07 | -5.09 | -0.016 |
| rs62054815 | SPPL2C | 1 | 0 | 5 | A | G | 0.23 | 3.5E-07 | -5.10 | -0.016 |
| rs12185233 | SPPL2C | 1 | 25.6 | 1f | C | G | 0.23 | 1.5E-07 | -5.26 | -0.017 |
| rs12373139 | SPPL2C | 1 | 0.53 | 1f | A | G | 0.23 | 1.4E-07 | -5.27 | -0.017 |
| rs63750417 | MAPT | 6 | 8.68 | 5 | T | C | 0.23 | 2.7E-07 | -5.14 | -0.017 |
| rs62063786 | MAPT | 6 | 7.65 | 5 | A | G | 0.23 | 2.3E-07 | -5.17 | -0.017 |
| rs17651549 | MAPT | 6 | 34 | 1f | T | C | 0.23 | 8.5E-08 | -5.36 | -0.017 |
| rs2526374 | RNF43 | 8 | 13.78 | 4 | T | G | 0.36 | 4.4E-08 | -5.47 | -0.015 |
| rs3744108 | MTMR4 | 6 | 23.1 | 5 | C | G | 0.38 | 7.7E-14 | -7.48 | -0.021 |
| rs6503870 | TEX14 | 20 | 9.53 | 7 | T | C | 0.38 | 3.9E-14 | 7.56 | 0.021 |
| rs2270951 | DCC | 22 | 16.7 | 6 | T | C | 0.46 | 1.1E-13 | -7.43 | -0.020 |
| rs8108738 | MAST3 | 22 | 13.96 | 4 | A | G | 0.47 | 5.9E-10 | 6.19 | 0.017 |
| rs882610 | ZNF446 | 7 | 7.69 | - | A | G | 0.27 | 1.6E-07 | 5.24 | 0.016 |
| rs3752109 | MZF1 | 3 | 12.56 | - | T | C | 0.28 | 8.4E-09 | 5.76 | 0.017 |
| rs11553387 | DDX27 | 6 | 16.86 | - | T | G | 0.22 | 2.1E-09 | 5.99 | 0.019 |
| rs1130146 | DDX27 | 19 | 26.2 | 5 | A | G | 0.39 | 1.7E-11 | -6.73 | -0.018 |
| rs6512577 | ZNFX1 | 14 | 0 | 5 | T | C | 0.22 | 1.8E-09 | 6.01 | 0.019 |
| rs12628603 | TRIOBP | 7 | 23.2 | 5 | A | G | 0.38 | 1.9E-06 | 4.77 | 0.013 |
| rs9610841 | TRIOBP | 7 | 22.8 | 5 | A | C | 0.46 | 1.1E-07 | 5.32 | 0.014 |
| rs8140207 | TRIOBP | 9 | 14.09 | 5 | T | G | 0.28 | 2.1E-07 | -5.19 | -0.015 |

Note: CADD: Combined Annotation Dependent Depletion score; RDB: Regulome DB score; MAF: minor allele frequency; Z: z-score from the GWAS meta-analysis; Effect size: magnitude of Z score association in standard deviation units. \*=SNP is an independent lead SNP. Genes containing multiple ExNS are in bold.

To link the associated genetic variants to genes, we applied three gene-mapping strategies as implemented in FUMA^12^ (**Online Methods**). *Positional* gene-mapping aligned SNPs to 514 genes by location, *eQTL (expression quantitative trait loci)* gene-mapping matched cis-eQTL SNPs to 709 genes whose expression levels they influence, and *chromatin interaction* mapping annotated SNPs to 226 genes based on three-dimensional DNA-DNA interactions between the SNPs’ genomic region and nearby or distant genes (**Figure 2; Supplementary Figure 7-8; Supplementary Table 10-12**). Of 882 total unique genes, 438 genes were implicated by at least two mapping strategies and 129 by all 3 (**Figure 3**). Of these, 15 genes are particularly notable as they are implicated via chromatin interactions between two independent genomic risk loci (**Supplementary Table 11**). *VAMP4* (locus 14), shows interactions in 6 tissue types including interactions with locus 15 in the left ventricle (**Figure 2a**). *SATB2* (locus 44) is linked by interaction in liver tissue to locus 43 (**Figure 2b**). *MEF2* (locus 82) shows interactions with locus 83 in 5 tissues (**Figure 2c**). *FBXL17* and *MAN2A1* are in two independent loci (87 and 88 respectively); they are mapped by eQTL associations and chromatin interactions between the two loci in the left ventricle (**Figure 2c**). Loci 102 and 103 show multiple interactions in one of 7 tissue types that are mapped to 8 genes encoding zinc finger proteins or histones (**Figure 2d**). *ELAVL2* (locus 130) interacts with locus 129 in the left ventricle and is also mapped by intra-locus interactions in other tissues (**Figure 2e**). *ATF4* (locus 212) is mapped by eQTLs in 3 tissue types and chromatin interactions in 7 tissue types, and interacts with locus 213 in the left ventricle (**Figure 2f**).

**Figure 2.**
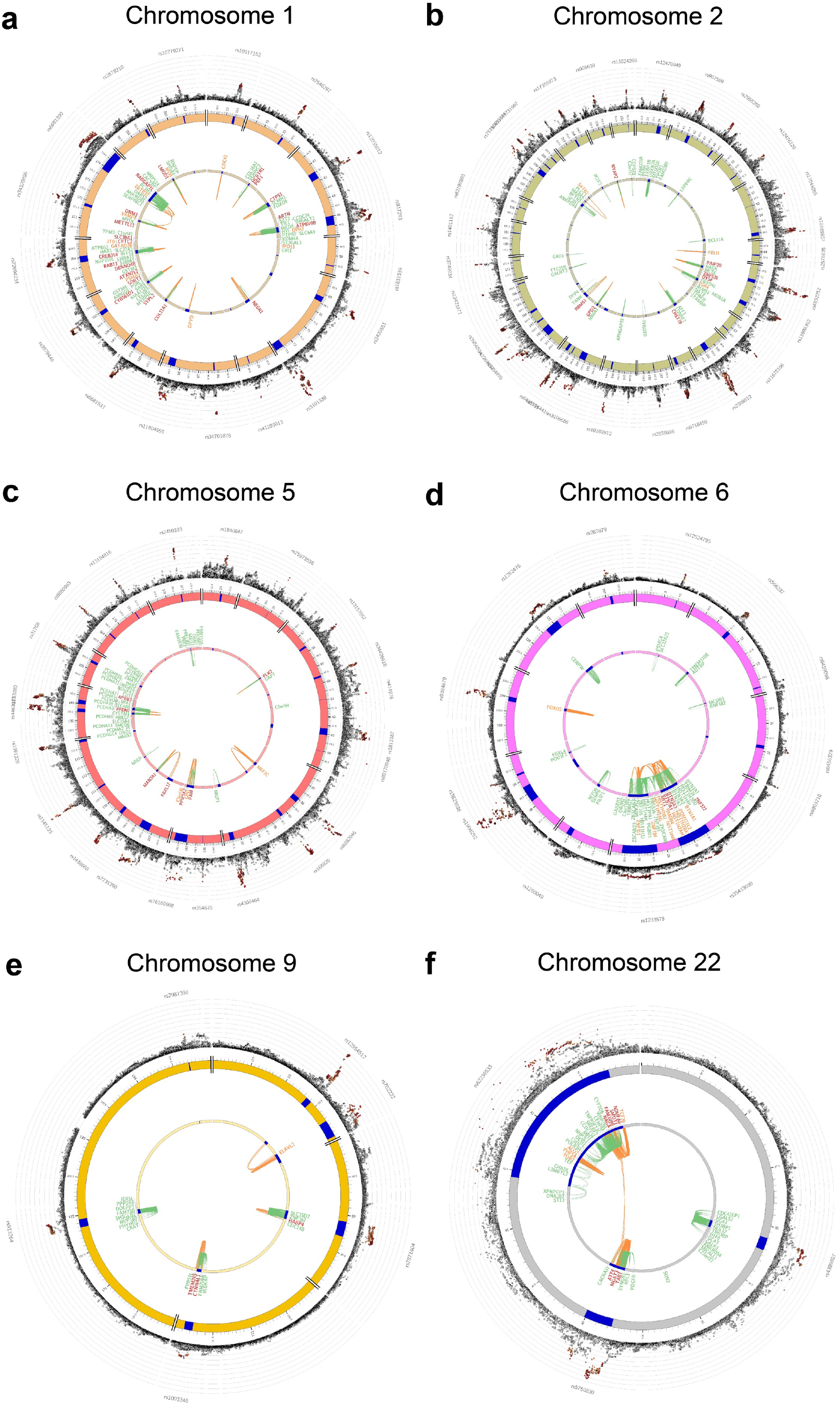
Genomic risk loci, expression quantitative trait locus (eQTL) associations and chromatin interactions for chromosomes containing cross-locus interactions. Circos plots showing genes on chromosomes 1 **(a)**, 2 **(b)** 5 **(c)** 6 **(d)** 9 **(e)** and 22 **(f)** that were implicated as genomic risk loci (blue regions) by positional mapping, eQTL mapping (green lines connecting an eQTL SNP to its associated gene), and/or chromatin interaction (orange lines connecting two interacting regions) and showed evidence of interaction across two independent genomic risk loci. Genes implicated by both eQTL and chromatin interactions mapping are in red. The outer layer shows a Manhattan plot containing the ─log10 transformed *P*-value of each SNP in the GWAS meta-analysis, with genome-wide significant SNPs in color corresponding to linkage disequilibrium patterns with the lead SNP. Circos plots for all chromosomes are provided in **Supplementary Fig. 7**.

**Figure 3.**
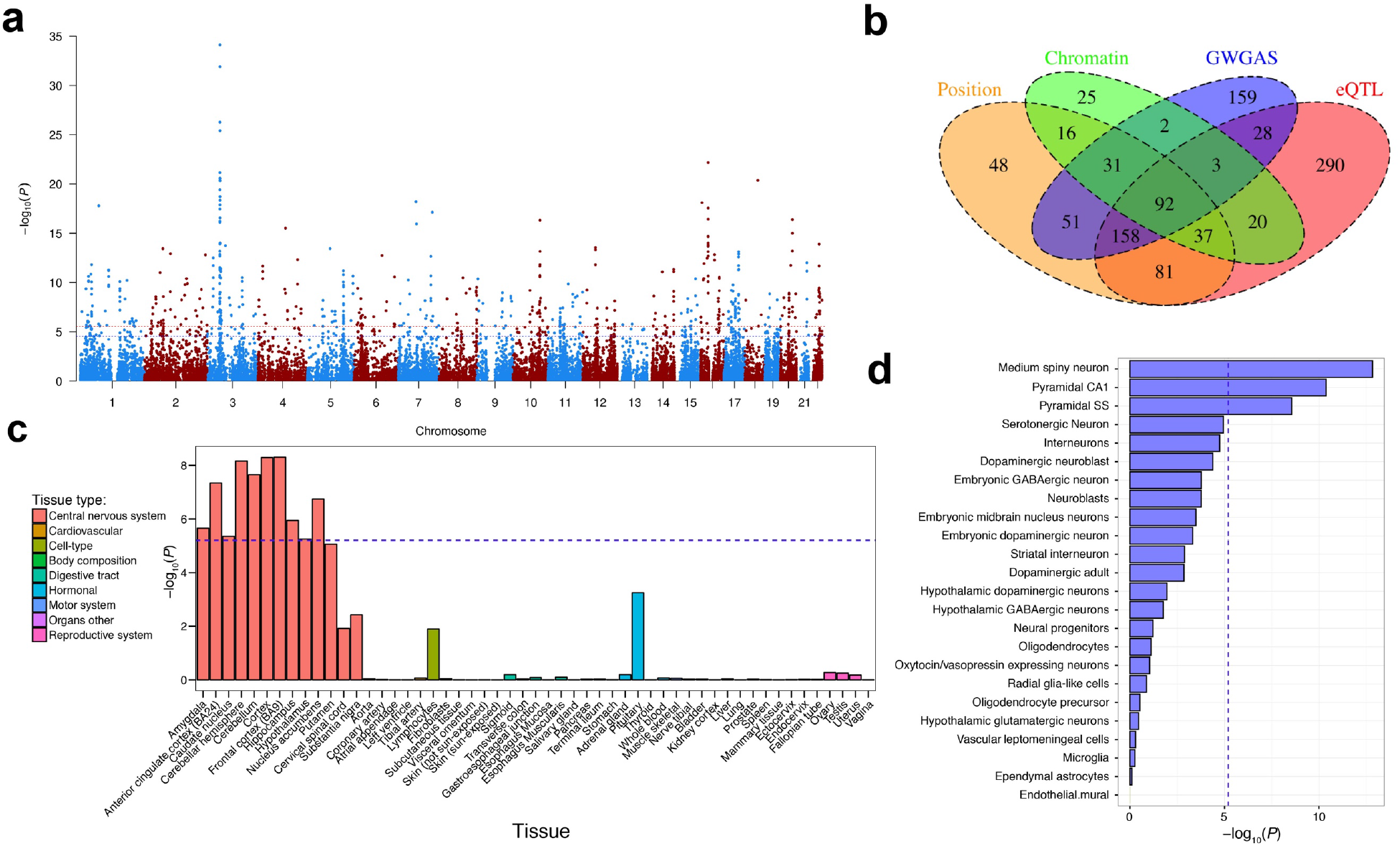
Mapping of genes and tissue- and cell expression profiles. (**a**) Manhattan plot of the genome-wide gene-based association analysis (GWGAS). The y-axis shows the ─log10 transformed *P*-value of each gene, and the chromosomal position on the x-axis. The red dotted line indicates the threshold for genome-wide significance of the gene-based test (*P*<2.76×10^−6^; 0.05/18,128), and the blue line indicates the suggestive threshold (*P*<2.76×10^−5^; 0.5/18,128) **(b)** Venn diagram showing overlap of genes implicated by positional mapping, eQTL mapping, chromatin interaction mapping, and GWGAS. **(c)** Gene expression profiles of identified genes for 53 tissue types. Expression data were extracted from the Genotype-Tissue Expression (GTEx) database. Expression values (RPKM) were log2 transformed with pseudocount 1 after winsorization at 50 and averaged per tissue. **(d)** Single-cell gene-expression analysis of genes related to intelligence in 24 cell-types. The dotted blue line indicates the Bonferroni-corrected significance threshold (*P*=0.05/7,323=6.83×10^−6^).

We performed genome-wide gene-based association analysis (GWGAS) using MAGMA^17^ (**Online Methods**). This approach provides aggregate association *P*-values based on all SNPs in a gene, whereas FUMA annotates individually significant SNPs to genes. GWGAS identified 524 associated genes (467 novel) (**Figure 3a; Supplementary Table 13; Supplementary Results 2.4.1**), of which 159 were outside of the GWAS risk loci, and 365 were also mapped by FUMA (**Figure 3b**). In total, 92 genes were implicated by all four strategies (**Supplementary Table 14**).

In gene-set analysis using the GWGAS results (**Online Methods**), six Gene Ontology^18^ gene-sets were significantly associated with intelligence: *neurogenesis* (Beta=0.153, SE=0.030, *P*=1.55×10^−7^), *neuron differentiation* (Beta=0.178, SE=0.038, *P*=1.36×10^−6^), *central nervous system neuron differentiation* (Beta=0.398, SE=0.089, *P*=3.97×10^−6^), *regulation of nervous system development* (Beta=0.187, SE=0.040, *P*=1.54×10^−6^), *positive regulation of nervous system development* (Beta=0.242, SE=0.052, *P*=1.93×10^−6^), and *regulation of synapse structure or activity* (Beta=0.153, SE=0.030, *P*=5.87×10^−6^) (**Supplementary Table 15**). Conditional analysis indicated that there were three independent associations, for the *neurogenesis, central nervous system neuron differentiation,* and *regulation of synapse structure or activity* processes, which together accounted for the associations of the other three sets (**Supplementary Results 2.4.2**).

Linking gene-based *P*-values to tissue-specific gene-sets (**Online Methods**), we observed strong associations across various brain areas (**Figure 3c; Supplementary Table 16; Supplementary Results 2.4.2**), most strongly with the cortex (*P*=5.12×10^−9^), and specifically frontal cortex (*P*=4.94×10^−9^). In brain single-cell expression gene-set analyses (Online Methods), we found significant associations of striatal medium spiny neurons (*P*=1.47×10^−13^) and pyramidal neurons in the CA1 hippocampal (*P*=4×10^−11^) and cortical somatosensory regions (*P*=3×10^−9^), (**Figure 3d; Supplementary Table 17**). Conditional analysis showed that the independent association signal in brain cells was driven by medium spiny neurons, neuroblasts, and pyramidal CA1 neurons (**Supplementary Results 2.4.2**).

Intelligence has been associated with a wide variety of human behaviors^19^ and brain anatomy^20^. Confirming previous reports^5,6^, we observed negative genetic correlations (Online methods) with ADHD (*r_g_*=-0.36, *P*=1.97×10^−24^), depressive symptoms (*r_g_*=-0.27, *P*=5.77×10^−10^), Alzheimer’s disease (*r_g_*=-0.26, *P*=5.77×10^−10^), and schizophrenia (*r_g_*=-0.22, *P*=2.58×10^−18^) and positive correlations with EA (*r_g_*=0.70, *P*<1×10^−200^) and longevity (*r_g_*=0.43, *P*=4.91×10^−8^) (**Supplementary Table 18; Supplementary Figure 9**). Comparison of our results with the contents of the NHGRI-21 EBI catalog^21^ supported these correlations, showing numerous shared genetic variants (**Supplementary Information 2.5; Supplementary Table 19-20**). Low enrichment (91 of 1,518 genes, hypergeometric *P*=0.03) was found for genes previously linked to intellectual disability or developmental delay (see **URLs; Online Methods**). However, our results replicate and add to previous genetic research on normal variation in intelligence, as catalogued in **Supplementary Tables 21-22**.

We used Mendelian Randomization (**Online Methods**) to test for potential credible causal associations between intelligence and genetically correlated traits (**Supplementary Table 23; Supplementary Figures 10-11**). We observed a strong effect of intelligence on EA (*bxy*=0.531, SE=0.006, *P*<1×10^−320^), that was bidirectional and showed a similar strong effect of EA on intelligence (*bxy*=0.517, SE=0.025, *P*=1.06×10^−96^), with only a small proportion of SNPs showing pleiotropic effects. Our result also suggested a protective effect of intelligence on ADHD (OR=0.46, *bxy*=-0.778, SE=0.051, *P*=3.80×10^−45^) and Alzheimer’s disease (OR=0.66, *bxy*=-0.411, SE=0.058, *P*=1.75×10^−12^). In line with a positive genetic correlation, we observed that intelligence was associated with higher risk of autism (OR=1.47, *bxy*=0.382, SE=0.099, *P*=1.10×10^−4^). There was evidence of a bidirectional association between intelligence and schizophrenia including a strong protective effect of intelligence on schizophrenia (OR=0.58, *bxy*=-0.551, SE=0.043, *P*=3.50×10^−30^), and a relatively smaller reverse effect (*bxy*= -0.195, SE= 0.012, *P*=2.02×10^−57^), with additional evidence for pleiotropy (**Supplementary Results 2.5.3**).

In conclusion, we conducted a large-scale genome-wide meta-analysis of intelligence in 279,930 individuals, resulting in the identification of 191 novel loci and 963 novel genes, and replicating previous associations with 15 loci and 78 genes. The applied combined strategies of functional annotation and gene-mapping and the use of unique biological data resources provide extensive information on functional consequences of relevant genetic variants and novel insight into underlying neurobiological pathways, and point towards the involvement of specific cell types. We also found suggestive evidence of causal associations between intelligence and neuropsychiatric traits. These results are important not only for understanding the biological underpinnings of individual differences in intelligence, but also contribute to our understanding of cognitive and related psychiatric disorders.

http://ukbiobank.ac.uk

http://www.biorxiv.org/content/early/2017/07/20/166298

http://hrsonline.isr.umich.edu

http://genesforgood.org

https://www.ncbi.nlm.nih.gov/gap

https://icar-project.com/

http://fuma.ctglab.nl

http://ctg.cncr.nl/software/magma

http://genome.sph.umich.edu/wiki/METAL_Program

https://github.com/bulik/ldsc

http://ldsc.broadinstitute.org/

https://data.broadinstitute.org/alkesgroup/LDSCORE/

http://www.genecards.org

http://www.med.unc.edu/pgc/results-and-downloads

http://software.broadinstitute.org/gsea/msigdb/collections.jsp

https://www.ebi.ac.uk/gwas/

https://github.com/ivankosmos/RegionAnnotator

http://cnsgenomics.com/software/gsmr/

## Acknowledgments

This work was funded by The Netherlands Organization for Scientific Research (NWO VICI 453-14-005 and NWO VIDI 452-12-014) and the Sophia Foundation for Scientific Research (grant nr: S14-27). The analyses were carried out on the Genetic Cluster Computer, which is financed by the Netherlands Scientific Organization (NWO: 480-05-003), by the VU University, Amsterdam, The Netherlands, and by the Dutch Brain Foundation, and is hosted by the Dutch National Computing and Networking Services SurfSARA. This research has been conducted using the UK Biobank resource under application number 16406 and the Database of Genotypes and Phenotypes (dbGaP) under accession numbers phs000280.v3.p1 and phs000209.v13.p3. We thank the numerous participants, researchers, and staff from many studies who collected and contributed to the data. Additional acknowledgments can be found in the Supplementary Information file. Summary statistics will be made available for download upon publication from http://ctglab.vu.nl.

## Author Contributions

D.P., J.E.S. and P.R.J. performed the analyses. D.P. conceived the idea of the study and supervised analyses. S.St. performed QC on the UK Biobank data and wrote the analysis pipeline. K.W. constructed and applied the FUMA pipeline for performing follow-up analyses. J.B. conducted the single cell enrichment analyses. C.L, M.N., A.R.H., T.J.C.P., and S.v.d.S. assisted with pipeline development and data analysis. S.A., P.B.B., J.R.I.C., K.L.G., J.K., R.K., E.K., M.L., M.N., C.A.R., J.W.T., H.Y., D.Z., S.H., N.K.H., I.K.K., S.L., G.W.M., A.B.M.-M., E.B.Q., G.S., N.S., B.T.W., T.W., D.E.A., D.K.A., D.A., R.M.B., P.B., K.E.B., T.D.C., O.C.-F., A.C., E.T.C., E.C., A.C., G.D., I.J.D., P.D., D.D., S.D., G.D., E.D.C., J.G.E., T.E., A.R.H., A.H., M.C.K., E.K., B.K., J.L., S.L., T.L., D.C.L., E.L., A.J.L., A.K.M., I.M., D.M., A.C.N., W.O., A.Palotie., A.Payton., N.P., R.A.P., K.R., I.R., P.R., D.R., F.W.S., M.A.S., O.B.S., N.S., J.M.S., V.M.S., N.C.S., R.E.S., K.S., A.N.V., D.R.W., E.W., J.Y., G.A., O.A.A., G.B., L.C., B.D., D.M.D., A.H., J.H.-L., M.A.I., K.S.K., N.G.M., S.E.M., N.L.P., R.P., T.J.C.P., S.R., P.F.S., H.T., S.I.V., and M.J.W. contributed data. D.P., J.E.S. and P.R.J. wrote the paper. All authors critically reviewed the paper.

## Author Information

PF Sullivan reports the following potentially competing financial interests: Lundbeck (advisory committee), Pfizer (Scientific Advisory Board member), and Roche (grant recipient, speaker reimbursement). G Breen reports consultancy and speaker fees from Eli Lilly and Illumina and grant funding from Eli Lilly. All other authors declare no financial interests or potential conflicts of interest.

Correspondence and requests for materials should be addressed to.

## Online methods

### Study Cohorts

The meta-analysis included new and previously reported GWAS summary statistics from 16 cohorts: UK Biobank (UKB), Cognitive Genomics Consortium (COGENT), Rotterdam Study (RS), Generation R Study (GENR), Swedish Twin Registry (STR), Spit for Science (S4S), High-IQ/Health and Retirement Study (HiQ/HRS), Twins Early Development Study (TEDS), Danish Twin Registry (DTR), IMAGEN, Brisbane Longitudinal Twin Study (BLTS), Netherlands Study of Cognition, Environment and Genes (NESCOG), Genes for Good (GfG), Swedish Twin Studies of Aging (STSA), Atherosclerosis Risk in Communities Study (ARIC), and the Multi-Ethnic Study of Atherosclerosis (MESA). Detailed descriptions of the samples, measures, genotyping, quality control, and analysis procedures for each cohort are provided in **Supplementary Information 1.1 and Supplementary**

### Meta-analysis

Stringent quality control measures were applied to the summary statistics for each GWAS cohort before combining. All files were checked for data integrity and accuracy. SNPs were filtered from further analysis if they met any of the following criteria: imputation quality (INFO/R^2^) score < 0.6, Hardy-Weinberg equilibrium (HWE) *P* < 5×10^−6^, study-specific minor allele frequency (MAF) corresponding to a minor allele count (MAC) < 100, and mismatch of alleles or allele frequency difference greater than 20% from the Haplotype Reference Consortium (HRC) genome reference panel^16^. Some cohorts used more stringent criteria (see **Supplementary Information 1.1**). Indels and SNPs that were duplicated, multi-allelic, monomorphic, or ambiguous (A/T or C/G) with a MAF >0.4 were also excluded. Visual inspection of the distribution of the summary statistics was completed, and Manhattan plots and QQ plots were created for the cleaned statistics from each cohort (**Supplementary Figure 1**).

The SNP association *P*-values from the GWAS cohorts were meta-analyzed with METAL^22^ (see **URLs**) in two phases. First, we meta-analyzed all cohorts with quantitative phenotypes (all except HiQ/HRS) using a sample-size weighted scheme. In the second phase, we added the HiQ/HRS study results to the first phase results, weighting each set of summary statistics by their respective non-centrality parameter (NCP). This method improves power when using an extreme case sampling design such as HiQ^23^. NCPs were estimated using the Genetic Power Calculator, as described by Coleman et al.^25^. After combining all data, meta-analysis results were further filtered to exclude any variants with N < 50,000.

The X chromosome was treated separately in the meta-analysis because imputed genotypes were not available for the X chromosome in the largest cohort (UKB), and there was little overlap between the UKB called genotypes and imputed data from other cohorts (NSNPs < 500). We therefore included only the called X chromosome variants in UKB for these analyses after performing X-specific quality control steps^26^.

We conducted a series of meta-analyses on subsets of the full sample using the same methods as above. Age group-specific meta-analyses were run in the cohorts of children (age < 17; GENR, TEDS, IMAGEN, BLTS; N=9,814), young adults (age ∼17-18; S4S, STR; N=6,033), and adults (age > 18, primarily middle-aged or older: UKB, RS, DTR, NESCOG, STSA, ARIC, MESA; N=214,291), excluding studies whose samples overlapped multiple age groups (COGENT, HiQ/HRS, GfG; N=49,792). To create independent discovery samples for use in polygenic score validation, we also conducted meta-analyses with a “leave-one-out” strategy in which summary statistics from four validation datasets were, respectively, excluded from the meta-analysis (see *Polygenic Scoring*, below).

### Cohort Heritability and Genetic Correlation

LD score regression^8^ was used to estimate genomic inflation and heritability of the intelligence phenotypes in each of the 16 cohorts using their post-quality control summary statistics, and to estimate the cross-cohort genetic correlations^27^. Pre-calculated LD scores from the 1000 Genomes European reference population were obtained from https://data.broadinstitute.org/alkesgroup/LDSCORE/. Genetic correlations were calculated on HapMap3 SNPs only. LD score regression was also used on the age subgroup meta-analyses to estimate heritability and cross-age genetic correlations.

### Genomic Risk Loci Definition

Independently associated loci from the meta-analysis were defined using FUMA^12^ (http://fuma.ctglab.nl/), an online platform for functional mapping of genetic variants. We first identified *independent significant SNPs* which have genome-wide significant *P*-value (<5×10^−8^) and represented signals that are independent from each other at *r^2^* <0.6. These SNPs were further represented by *lead SNPs*, which are a subset of the independent significant SNPs that are in approximate linkage equilibrium with each other at *r^2^* <0.1. We then defined associated *genomic risk loci* by merging any physically overlapping lead SNPs (linkage disequilibrium [LD] blocks <250kb apart). Borders of the genomic risk loci were defined by identifying all SNPs in LD (*r*^*2*^≧0.6) with one of the independent significant SNPs in the locus, and the region containing all of these *candidate SNPs* was considered to be a single independent genomic risk locus. All LD information was calculated from UK Biobank genotype data.

### Proxy-replication with Educational Attainment (EA)

We conducted GWAS of EA, an outcome with a high genetic correlation with intelligence^5^, in a non-overlapping European subset of the UKB sample (N=188,435) who did not complete the intelligence measure. EA was coded as maximum years of education completed, using the same methods as earlier analyses^28^ and GWAS was conducted using the same quality control and analytic procedures as described for the UKB intelligence phenotype (**Supplementary Information 1.1**). To test replication of the SNPs with this proxy phenotype, we performed a sign concordance test for all GWS SNPs from the meta-analysis using the exact binomial test. For each independent genomic locus, we considered it to be evidence for replication if the lead SNP or another correlated SNP in the region was sign concordant with the corresponding SNP in the intelligence meta-analysis and had a *P*-value of association with EA smaller than 0.05/246=0.0002.

### Polygenic Scoring

We calculated polygenic scores (PGS) based on the SNP effect sizes of the leave-one-out metaanalyses, from which four cohorts were (separately) excluded and reserved for score validation. These included a child (GENR), young adult (S4S), and adult sample (RS). We also included the UKB-wb sample to test for validation in a very large (N = 53,576) cohort with the greatest phenotypic similarity to the largest contributor to the meta-analysis statistics (UKB-ts), in order to maximize potential predictive power. PGS were calculated on the genotype data using LDpred^10^, a Bayesian PGS method that utilizes a prior on effect size distribution to remodel the SNP effect size and account for LD, and PRSice^9^, a PLINK^29^-based program that automates optimization of the set of SNPs included in the PGS based on a high-resolution filtering of the GWAS *P*-value threshold. LDpred PGS were applied to the called, cleaned, genotyped variants in each of the validation cohorts with UK Biobank as the LD reference panel. PRSice PGS were calculated on hard-called imputed genotypes using *P-*value thresholds from 0.0 to 0.5 in steps of 0.001. The explained variance (Δ*R*^2^) was derived from a linear model in which the GWAS intelligence phenotype was regressed on each PGS while controlling for the same covariates as in each cohort-specific GWAS, compared to a linear model with GWAS covariates only.

### Stratified Heritability

We partitioned SNP heritability using stratified LD Score regression^30^ in three ways: 1) by functional annotation category, 2) by minor allele frequency (MAF) in six percentile bins, and 3) by chromosome. Annotations for 22 binary categories of functional genomic characteristics (e.g. coding or regulatory regions) were obtained from the LD score website (https://github.com/bulik/ldsc). The Bonferroni-corrected significance threshold was .05/50 annotations=.001.

### Functional Annotation of SNPs

Functional annotation of SNPs implicated in the meta-analysis was performed using FUMA^12^ (http://fuma.ctglab.nl/). We selected all *candidate SNPs* in genomic risk loci having an *r*^*2*^≧0.6 with one of the independent significant SNPs (see above), a *P*-value (*P*<1e-5) and a MAF>0.0001 for annotations. Functional consequences for these SNPs were obtained by matching SNPs’ chromosome, base-pair position, and reference and alternate alleles to databases containing known functional annotations, including ANNOVAR^31^ categories, Combined Annotation Dependent Depletion (CADD) scores^13^, RegulomeDB^14^ (RDB) scores, and chromatin states^15,16^. ANNOVAR categories identify the SNP’s genic position (e.g. intron, exon, intergenic) and associated function. CADD scores predict how deleterious the effect of a SNP is likely to be for a protein structure/function, with higher scores referring to higher deleteriousness. A CADD score above 12.37 is the threshold to be potentially pathogenic^13^. The RegulomeDB score is a categorical score based on information from expression quantitative trait loci (eQTLs) and chromatin marks, ranging from 1a to 7 with lower scores indicating an increased likelihood of having a regulatory function. Scores are as follows: 1a=eQTL + Transciption Factor (TF) binding + matched TF motif + matched DNase Footprint + DNase peak; 1b=eQTL + TF binding + any motif + DNase Footprint + DNase peak; 1c=eQTL + TF binding + matched TF motif + DNase peak; 1d=eQTL + TF binding + any motif + DNase peak; 1e=eQTL + TF binding + matched TF motif; 1f=eQTL + TF binding / DNase peak; 2a=TF binding + matched TF motif + matched DNase Footprint + DNase peak; 2b=TF binding + any motif + DNase Footprint + DNase peak; 2c=TF binding + matched TF motif + DNase peak; 3a=TF binding + any motif + DNase peak; 3b=TF binding + matched TF motif; 4=TF binding + DNase peak; 5=TF binding or DNase peak; 6=other;7=Not available. The chromatin state represents the accessibility of genomic regions (every 200bp) with 15 categorical states predicted by a hidden Markov model based on 5 chromatin marks for 127 epigenomes in the Roadmap Epigenomics Project^16^. A lower state indicates higher accessibility, with states 1-7 referring to open chromatin states. We annotated the minimum chromatin state across tissues to SNPs. The 15-core chromatin states as suggested by Roadmap are as follows: 1=Active Transcription Start Site (TSS); 2=Flanking Active TSS; 3=Transcription at gene 5’ and 3’; 4=Strong transcription; 5= Weak Transcription; 6=Genic enhancers; 7=Enhancers; 8=Zinc finger genes & repeats; 9=Heterochromatic; 10=Bivalent/Poised TSS; 11=Flanking Bivalent/Poised TSS/Enh; 12=Bivalent Enhancer; 13=Repressed PolyComb; 14=Weak Repressed PolyComb; 15=Quiescent/Low. Standardized SNP effect sizes were calculated for the most impactful SNPs by transforming the sample size-weighted meta-analysis *Z* score, as described in Zhu et al., 2016^32^.

### Gene-mapping

Genome-wide significant loci obtained by GWAS were mapped to genes in FUMA^12^ using three strategies:

1. Positional mapping maps SNPs to genes based on physical distance (within a 10kb window) from known protein coding genes in the human reference assembly (GRCh37/hg19).
2. eQTL mapping maps SNPs to genes with which they show a significant eQTL association (i.e. allelic variation at the SNP is associated with the expression level of that gene). eQTL mapping uses information from 45 tissue types in 3 data repositories (GTEx^33^, Blood eQTL browser^34^, BIOS QTL browser^35^), and is based on cis-eQTLs which can map SNPs to genes up to 1Mb apart. We used a false discovery rate (FDR) of 0.05 to define significant eQTL associations.
3. Chromatin interaction mapping was performed to map SNPs to genes when there is a three-dimensional DNA-DNA interaction between the SNP region and another gene region. Chromatin interaction mapping can involve long-range interactions as it does not have a distance boundary. FUMA currently contains Hi-C data of 14 tissue types from the study of Schmitt et al^36^. Since chromatin interactions are often defined in a certain resolution, such as 40kb, an interacting region can span multiple genes. If a SNPs is located in a region that interacts with a region containing multiple genes, it will be mapped to each of those genes. To further prioritize candidate genes, we selected only interaction-mapped genes in which one region involved in the interaction overlaps with a predicted enhancer region in any of the 111 tissue/cell types from the Roadmap Epigenomics Project^16^ and the other region is located in a gene promoter region (250bp up and 500bp downstream of the transcription start site and also predicted by Roadmap to be a promoter region). This method reduces the number of genes mapped but increases the likelihood that those identified will have a plausible biological function. We used a FDR of 1×10^−5^ to define significant interactions, based on previous recommendations^36^ modified to account for the differences in cell lines used here.

### Functional annotation of mapped genes

Genes implicated by mapping of significant GWAS SNPs were further investigated using the GENE2FUNC procedure in FUMA^12^, which provides hypergeometric tests of enrichment of the list of mapped genes in 53 GTEx^33^ tissue-specific gene expression sets, 7,246 MSigDB gene-sets^37^, and 2,195 GWAS catalog gene-sets^21^. The Bonferroni-corrected significance threshold was 0.05/9,494 gene-sets=5.27×10^−6^.

### Gene-based analysis

SNP-based *P-*values from the meta-analysis were used as input for the gene-based genome-wide association analysis (GWGAS). 18,128 protein-coding genes (each containing at least 1 GWAS SNP) from the NCBI 37.3 gene definitions were used as basis for GWGAS in MAGMA (http://ctg.cncr.nl/software/magma)^17^. The Bonferroni-corrected genome-wide significance threshold was .05/18,128 genes=2.76×10^−6^.

### Gene-set analysis

Results from the GWGAS analyses were used to test for association in three types of predefined gene-sets:

1. 7,246 curated gene-sets representing known biological and metabolic pathways were derived from 9 data resources, catalogued by and obtained from the MsigDB version 5.2^29^ (http://software.broadinstitute.org/gsea/msigdb/collections.jsp)
2. gene expression values from 53 tissues obtained from GTEx^33^, log2 transformed with pseudocount 1 after winsorization at 50 and averaged per tissue
3. cell-type specific expression in 24 types of brain cells, which were calculated following the method described in Skene et al^38^. and Coleman et al^25^. Briefly, brain cell-type expression data was drawn from single-cell RNA sequencing data from mouse brains. For each gene, the value for each cell-type was calculated by dividing the mean Unique Molecular Identifier (UMI) counts for the given cell type by the summed mean UMI counts across all cell types. Single-cell gene-sets were derived by grouping genes into 40 equal bins by specificity of expression.

These gene-sets were tested using MAGMA. We computed competitive *P*-values, which represent the test of association for a specific gene-set compared to other gene-sets. This method is more robust to Type I error than self-contained tests that only test for association of a gene-set against the null hypothesis of no association^39^. The Bonferroni-corrected significance threshold was 0.05/7,323 gene-sets=6.83×10^−6^. Conditional analyses were performed as a followup using MAGMA to test whether each significant association observed was independent of all others. The association between each gene-set was tested conditional on the most strongly associated set, and then - if any substantial (p<.05/number of gene-sets) associations remained - by conditioning on the first and second most strongly associated set, and so on until no associations remained. Gene-sets that retained their association after correcting for other sets were considered to be independent signals. We note that this is not a test of association per se, but rather a strategy to identify, among gene-sets with known significant associations whose defining genes may overlap, which set(s) are responsible for driving the observed association.

### Cross-Trait Genetic Correlation

Genetic correlations (*r_g_*) between intelligence and 38 phenotypes were computed using LD score regression^27^, as described above, based on GWAS summary statistics obtained from publicly available databases (http://www.med.unc.edu/pgc/results-and-downloads; http://ldsc.broadinstitute.org/; **Supplementary Table 18**). The Bonferroni-corrected significance threshold was 0.05/38 traits=1.32×10^−3^.

### GWAS catalog lookup

We used FUMA to identify SNPs with previously reported (*P* < 5×10^−5^) phenotypic associations in published GWAS listed in the NHGRI-EBI catalog^21^ which overlapped with the genomic risk loci identified in the meta-analysis. As an additional relevant phenotype of interest, we examined whether the genes associated with intelligence in this study (by FUMA mapping or GWGAS) were overrepresented in a set of 1,518 genes linked to intellectual disability and/or developmental delay, as compiled by RegionAnnotater (https://github.com/ivankosmos/RegionAnnotator). Many of these have been identified by non-GWAS sources and are not represented in the NHGRI catalog. We tested for enrichment using a hypergeometric test with a background set of 19,283 genomic protein-coding genes, as in FUMA. Manual lookups were also performed to identify overlapping loci/genes with known previous GWAS of intelligence.

### Mendelian Randomization

To infer credible causal associations between intelligence and traits that are genetically correlated with intelligence, we performed Generalised Summary-data based Mendelian Randomization^40^ (GSMR; http://cnsgenomics.com/software/gsmr/). This method utilizes summary-level data to test for causal associations (*b_xy_*) between a risk factor and an outcome by using genome-wide significant SNPs as instrumental variables. HEIDI-outlier detection was used to filter genetic instruments that show clear pleiotropic effects on the exposure phenotype and the outcome phenotype. We used a threshold p-value of 0.01 for the outlier detection analysis in HEIDI which removes 1% of SNPs by chance if there is no pleiotropic effect. To test for a causal effect of intelligence (*b_zx_*) on an outcome (*b_zy_*) we selected traits in non-overlapping samples that showed significant genetic correlations (*r_g_*) with intelligence. We tested for bi-directional causation by repeating the analyses using independent GWS SNPs related to the outcome phenotypes as exposure and intelligence as the outcome phenotype. For each trait, we selected independent (*r*^*2*^=<0.1), GWS lead SNPs as instrumental variables in the analyses. For traits with less than 10 lead SNPs (i.e. the minimum number of SNPs on which GSMR can perform a reliable analysis) we selected independent SNPs (*r*^*2*^=<0.1), with a GWS *P*-value (<5×10^−8^), except for ADHD for which the threshold was lowered to 1×10^−5^ due to the small number of GWS SNPs. The estimated *b_zx_* and *b_zy_* are approximately equal to the natural log odds ratio (OR)^40^. An OR of 2 can be interpreted as a doubled risk compared to the population prevalence of a binary trait for every SD increase in the exposure trait. For quantitative traits the *b_zx_* and *b_zy_* can be interpreted as a one standard deviation increase explained in the outcome trait for every SD increase in the exposure trait.

### Data availability

Summary statistics will be made available for download upon publication (https://ctg.cncr.nl).

